# SARS-CoV-2 NSP13 interacts with TEAD to suppress Hippo-YAP signaling

**DOI:** 10.1101/2023.11.30.569413

**Authors:** Fansen Meng, Jong Hwan Kim, Chang-Ru Tsai, Jeffrey D. Steimle, Jun Wang, Yufeng Shi, Rich G. Li, Bing Xie, Vaibhav Deshmukh, Shijie Liu, Xiao Li, James F. Martin

**Affiliations:** McGill Gene Editing Lab, The Texas Heart Institute, Houston, Texas, United States; Cardiomyocyte Renewal Laboratory, The Texas Heart Institute, Houston, Texas, United States; Department of Integrative Physiology, Baylor College of Medicine, Houston, Texas, United States; The Heart Institute, Division of Molecular Cardiovascular Biology, Cincinnati Children’s Hospital Medical Center, Cincinnati, OH, United States; Department of Pediatrics, University of Cincinnati College of Medicine, Cincinnati, OH, United States

**Keywords:** NSP13, YAP, TEAD4, SARS-CoV-2, Helicase, Hippo pathway, TTF2

## Abstract

The Hippo pathway controls organ development, homeostasis, and regeneration primarily by modulating YAP/TEAD-mediated gene expression. Although emerging studies report Hippo-YAP dysfunction after viral infection, it is largely unknown in the context of severe acute respiratory syndrome coronavirus 2 (SARS-CoV-2). Here, we analyzed RNA sequencing data from induced pluripotent stem cell–derived cardiomyocytes (iPSC-CMs) and SARS-CoV-2-infected human lung samples, and observed a decrease in YAP target gene expression. In screening SARS-CoV-2 nonstructural proteins, we found that nonstructural protein 13 (NSP13), a conserved coronavirus helicase, inhibits YAP transcriptional activity independent of the upstream Hippo kinases LATS1/2. Consistently, introducing NSP13 into cardiomyocytes suppresses an active form of YAP (YAP5SA) *in vivo*. Subsequent investigations on NSP13 mutants revealed that NSP13 helicase activity, including DNA binding and unwinding, is crucial for suppressing YAP transactivation. Mechanistically, TEAD4 serves as a platform to recruit NSP13 and YAP. NSP13 likely inactivates the YAP/TEAD4 transcription complex by remodeling chromatin to recruit proteins, such as transcription termination factor 2 (TTF2), to bind the YAP/TEAD/NSP13 complex. These findings reveal a novel YAP/TEAD regulatory mechanism and uncover molecular insights into Hippo-YAP regulation after SARS-CoV-2 infection.

## Introduction

The evolutionarily conserved Hippo-YAP signaling pathway integrates various extracellular signals, including mechanical force, cell adhesion, and nutrient availability, through the protein kinases LATS1/2 (*1*). YAP is the primary substrate of LATS kinases, and following phosphorylation, is inactivated via cytoplasmic retention or degradation. Unphosphorylated YAP translocates to the nucleus and complexes with transcription factor partners, notably TEAD, to activate target genes (*1, 2*) involved in tissue development, homeostasis, and regeneration across multiple organs (*2–7*). YAP/TEAD is regulated by intricate molecular mechanisms that govern its activity, localization, and interaction. Dysregulation of these processes has been implicated in various diseases, including cancer (*8–11*).

Recent studies reveal that YAP is an endogenous brake of innate antiviral immunity (*12, 13*). Genetic loss of YAP enhances antiviral responses, whereas expression of a transactivation deficient mutant (YAP 6SA) suppresses these responses. Conversely, multiple canonical YAP target genes, such as *IL-6*, *Ccl2*, and *Csf1*, modulate innate immune signaling and cytokine production (*14–16*). Notably, different types of viral infection distinctly influence YAP expression, degradation, and nuclear localization (*17*). Although these observations shed light on the complex dynamics of YAP during viral responses, the mechanisms underlying YAP/TEAD regulation after SARS-CoV-2 infection are poorly understood. SARS-CoV-2, the center of the COVID-19 pandemic, has well-documented effects on the respiratory, digestive, central nervous, and cardiovascular systems (*18–20*). Investigating SARS-CoV-2 and Hippo-YAP signaling provides molecular insight into virus–induced pathophysiology. Two groups recently reported opposite effects on YAP activity after SARS-CoV-2 infection (*21, 22*). However, the lack of YAP/TEAD target gene expression data in these studies prevents a full understanding of these disparate results. Thus, the precise understanding of YAP/TEAD regulation after SARS-CoV-2 infection remains elusive.

Here, we found that SARS-CoV-2 infection reduces YAP target gene expression in human induced pluripotent stem cell–derived cardiomyocytes (hiPSC-CMs) and lung epithelial cells of COVID-19 patients. By comprehensively screening SARS-CoV-2 nonstructural proteins, we identified NSP13 as a key factor in inhibiting YAP transcriptional activity and suppressing active YAP5SA activity *in vivo*. Moreover, NSP13 suppression of YAP is dependent on its helicase activity, where both DNA binding and unwinding are required. Mechanistically, NSP13 is directly bound to TEAD4 and inhibits the transcriptional activity of the YAP/TEAD4 complex by remodeling chromatin and recruiting transcriptional suppressors such as TTF2. These findings reveal a novel function of NSP13 in regulating YAP/TEAD activity and provide key insights into how the SARS-CoV-2 genome modulates transcriptional activity of the YAP-TEAD complex.

## Results

### SARS-CoV-2 infection suppresses YAP activity in host cells

To assess YAP activity following SARS-CoV-2 infection *in vitro*, we evaluated a bulk RNA-sequencing dataset derived from hiPSC-CMs exposed to varying SARS-CoV-2 concentrations (*23*) (Figure 1A). Cardiomyocyte-specific YAP target gene (*24*) expression revealed that YAP targets were decreased in a dose-dependent manner after SARS-CoV-2 infection (Figure 1B). YAP is known to directly regulate several innate immunity genes, including *Ccl2*, *Thbs1*, and *Csf1* (*15, 16*). We observed that these genes are down-regulated in SARS-CoV-2 infected cells, mirroring the decrease in canonical YAP targets such as *Vgll3*, *Amotl2*, and *Ccn1* (Figure 1C).

**Figure 1.**
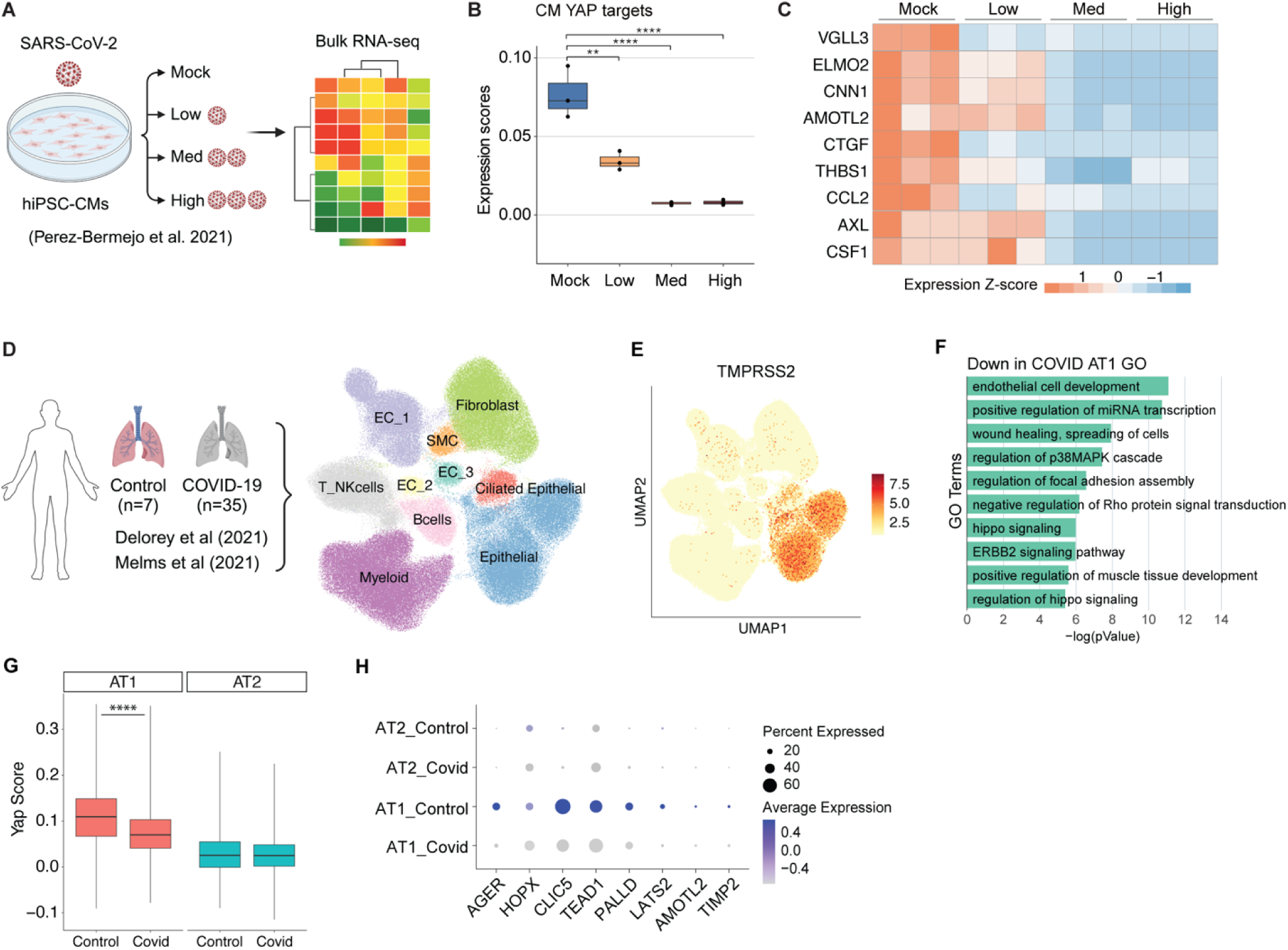
SARS-CoV-2 infection suppresses YAP activity. (**A**) Overview of SARS-CoV-2 infection in human induced pluripotent stem cell–derived cardiomyocytes (hiPSC-CMs). (**B**) Box plot showing the mean expression scores of known YAP target genes in hiPSC-CM bulk RNA-sequencing data. Each dot represents a biological replicate. Student *t-*test;**p< 0.01, ****p< 0.0001. (**C**) Heatmap displaying the expression Z-scores of example YAP targets in the iPSC-CM bulk RNA-sequencing data. Each column corresponds to a single biological sample. (**D**). Overview of integrated single-nucleus RNA sequencing and uniform manifold approximation and projection (UMAP) of cell types in lung samples from controls and patients with COVID-19. EC, endothelial cells; NK, natural killer cells; SMC, smooth muscle cells. (**E**) UMAP visualization of TMPRESS2 expression in the 10 cell types. **(F)** GO analysis of downregulated genes in AT1 cells from COVID patients. **(G)** Yap scores in alveolar type 1 (AT1) and alveolar type 2 (AT2) epithelial cells from lung samples in controls and patients with COVID-19. Wilcoxon test, ****p< 0.0001. **(H)** Expression of Yap targets in AT1 and AT2 cells of control and COVID-positive lung samples.

We then assessed YAP activity with SARS-CoV-2 infection *in vivo* by integrating and reanalyzing single-nuclei RNA (snRNA) sequencing data derived from human lung samples (*25, 26*). We identified 10 major cell types (Figure 1D, Figure Supplement 1A). TMPRSS2 and ACE2, two key entry factors for SARS-CoV-2 infection (*27*) were more highly expressed in lung epithelial cells compared to other cell types (Figure 1E, Figure Supplement 1B). Lung epithelial cells were categorized into alveolar type 1 (AT1) and alveolar type 2 (AT2) (Figure Supplement 1C, D). SARS-CoV-2 is more likely to infect AT1 cells as they cover >95% of the alveolar surface. We performed an unbiased GO analysis of the differentially expressed genes (DEGs) and observed that genes associated with or regulating the Hippo pathway were downregulated in AT1 cells from COVID-19 patients (Figure 1F). Consistent with previous data *in vitro,* the YAP score, evaluated via YAP target gene expression, was lower in AT1 cells from COVID-19 patients than in controls (Figure 1G). Signature genes associated with AT1 cells, including the reported YAP target genes *AGER* and *CLIC5* (*28, 29*) were downregulated in COVID-19 patient lung samples (Figure 1H). These data support the notion that SARS-CoV-2 infection suppresses YAP activity in host cells. However, further analysis revealed that multiple YAP target genes involved in innate immunity and cytokine signaling were paradoxically elevated (Figure Supplement 1E). This discrepancy likely reflects a combination of factors, including cell type specificity, activation of parallel signaling pathways, and alterations in mechanical cues or tissue architecture, that can independently drive expression of these genes.

### NSP13 inhibits YAP5SA transactivation

To identify which SARS-CoV-2 protein suppresses YAP activity, we screened 11 NSP proteins using a dual-luciferase reporter assay. This assay measures firefly and Renilla luciferase activities sequentially from the same sample. Normalization to Renilla luciferase serves as an internal control, ensuring accuracy and reproducibility of the results. Compared with other NSPs, NSP13 strongly inhibited YAP transcription activity (Figure 2A). By using a constitutively active form of YAP (YAP5SA) that is resistant to phosphorylation and inactivation by LATS1/2 (*30*), we observed that NSP13 suppresses YAP transactivation independent of its upstream kinase LATS2 (Figure Supplement 2A). Moreover, NSP13 attenuated YAP5SA activity dose-dependently (Figure 2B). A recent study reported that NSP13 suppresses episomal DNA transcription, as evidenced by reduced Renilla luciferase activity and decreased GFP expression upon co-expression with NSP13 (*31*). To evaluate this in our system, we examined our raw Renilla luciferase data and found that while 100 ng of NSP13 had no effect, 400 ng of NSP13 reduced Renilla luciferase levels by approximately 50% compared to the YAP5SA-only group (Figure Supplement 2B and 3C–D). Notably, the firefly luciferase signal, driven by YAP/TEAD interaction (HOP-Flash), exhibited an even greater reduction. To assess the specificity of this suppression, we also conducted a Notch reporter assay and observed that co-expression of NSP13 with NICD (Notch intracellular domain) did not inhibit Notch signaling (Figure Supplement 2C). These results reveal a specific suppressive effect of NSP13 on YAP-mediated transcriptional activity.

**Figure 2.**
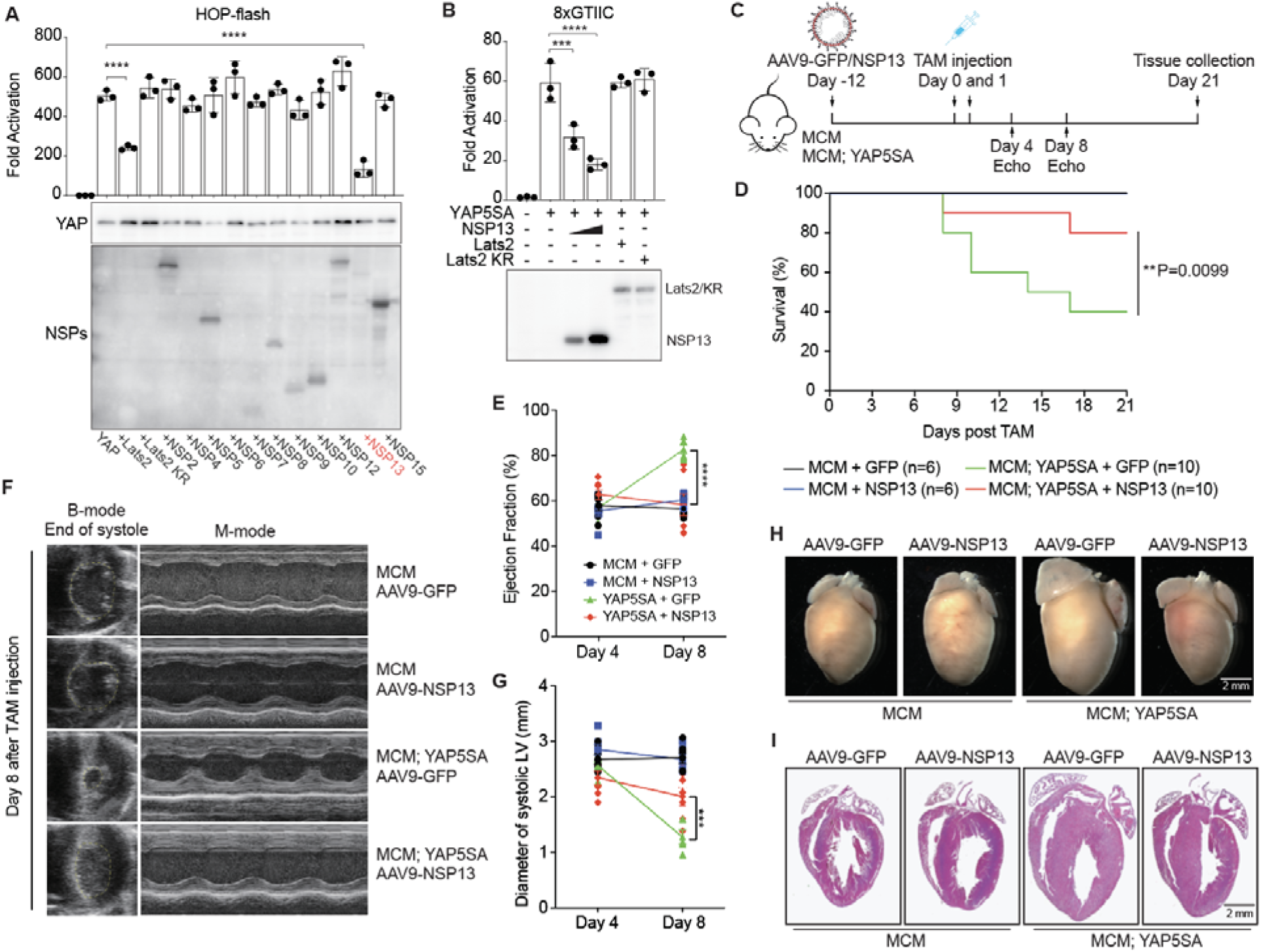
NSP13 inhibits YAP transactivation. (**A**) Screening of 11 NSPs for YAP activation by using a dual-luciferase reporter assay (HOP-flash). Compared with other NSPs, NSP13 strongly inhibited YAP transactivation at low protein expression levels. (n = 3 independent experiments; data are reported as mean ± SD). ****p< 0.0001, one-way ANOVA. (**B**) Reporter assay (8xGTIIC) results showing that NSP13 but not YAP upstream kinase LATS2 inhibited YAP5SA transactivation in a dose-dependent manner. (n = 3 independent experiments; data are presented as the mean ± SD). ***p< 0.001, ****p< 0.0001, one-way ANOVA. (**C**) Experimental design of NSP13 study in mice. Control (aMyHC-MerCreMer;WT) and YAP5SA (aMyHC-MerCreMer;YAP 5SA) mice were injected with AAV9-GFP or AAV9-NSP13. At 12 days after virus injection, the mice received two low doses of TAM (10 ug/g). Cardiac function was recorded by echocardiography at day 4 and 8 after the second shot of tamoxifen. Hearts of all surviving mice were collected at day 21 post-tamoxifen injection. (**D**) NSP13 expression in cardiomyocytes improved the survival rate of YAP5SA mice after TAM injection compared to YAP5SA mice with AAV9-GFP infection. **p=0.0099, log-rank (Mantel-Cox) test. (**E**) Ejection fraction in YAP5SA mice was increased on day 8 after tamoxifen injections (10 ug/g x2). NSP13 expression reversed the increase of EF in YAP5SA mice. ****p<0.0001, three-way ANOVA. (**F**) Representative B-mode and M-mode echocardiographic images of mouse hearts in four groups, 8 days after tamoxifen (TAM) induction. (**G**) A reduction in the left ventricle size was seen in YAP5SA mice at day 8 after tamoxifen injection. NSP13 introduction reversed this trend as evidenced by an increase in the diameter of the left ventricle. ***p< 0.001, three-way ANOVA. (**H-I**) Representative whole mount and hematoxylin & eosin images of mouse hearts at 21 days after tamoxifen induction. Scale bar, 2 mm.

To investigate NSP13 function *in vivo*, we used YAP5SA transgenic mice (aMyHC-MerCreMer;YAP5SA), which express YAP5SA in cardiomyocytes following tamoxifen administration. YAP5SA induction produces cardiomyocyte hyperplasia and increased ejection fraction (EF) and fractional shortening (FS), ultimately leading to mortality (*24*). We used the adenovirus expressing NSP13 (AAV9-NSP13), which specifically infects cardiomyocytes (Figure Supplement 2D and Figure 2C). NSP13-induced expression in YAP5SA mouse cardiomyocytes increased survival rates (Figure 2D) and restored cardiac function (Figure 2E and Figure Supplement 2E). Furthermore, NSP13 expression reversed the smaller left ventricle (LV) chamber in YAP5SA mouse hearts (*24*) (Figure 2F, G). To investigate NSP13 induction in the mouse heart further, we collected heart tissue from surviving mice 21 days after tamoxifen injection. Notably, NSP13 expression in YAP5SA mice reversed heart overgrowth (Figure 2H, I, and Figure Supplement 2F). Together, these data indicate that NSP13 suppresses YAP activity *in vitro* and *in vivo*.

### NSP13 helicase activity is required for YAP suppression

NSP13, a helicase with conserved sequence across all coronaviruses (Figure 3A), plays a critical role in viral replication and is a promising target for antiviral treatment (*32–39*). K131, K345/K347, and R567 were identified as important amino acid sites for NSP13 helicase activity in SARS-CoV (*40*). Given the 99.8% sequence identity of NSP13 between SARS-CoV-2 and SARS-CoV, we speculated that these sites were similarly crucial for SARS-CoV-2 NSP13. Reporter assays revealed that NSP13-K131A, which retains partial helicase activity, can suppress YAP, whereas NSP13-R567A (no ATP consumption) and NSP13-K345A/K347A (obstructed nucleic acid binding channel) failed to inhibit YAP transcriptional activity (Figure 3B). To assess the contribution of each domain to YAP regulation, we constructed six NSP13 truncations (Figure 3C), yet none of these truncations reduced YAP transactivation (Figure 3D). These findings suggest that full-length NSP13 with helicase activity (DNA binding and ATP dependent unwinding) is required to suppress YAP transactivation, and that partial helicase function is sufficient to fully inhibit YAP.

**Figure 3.**
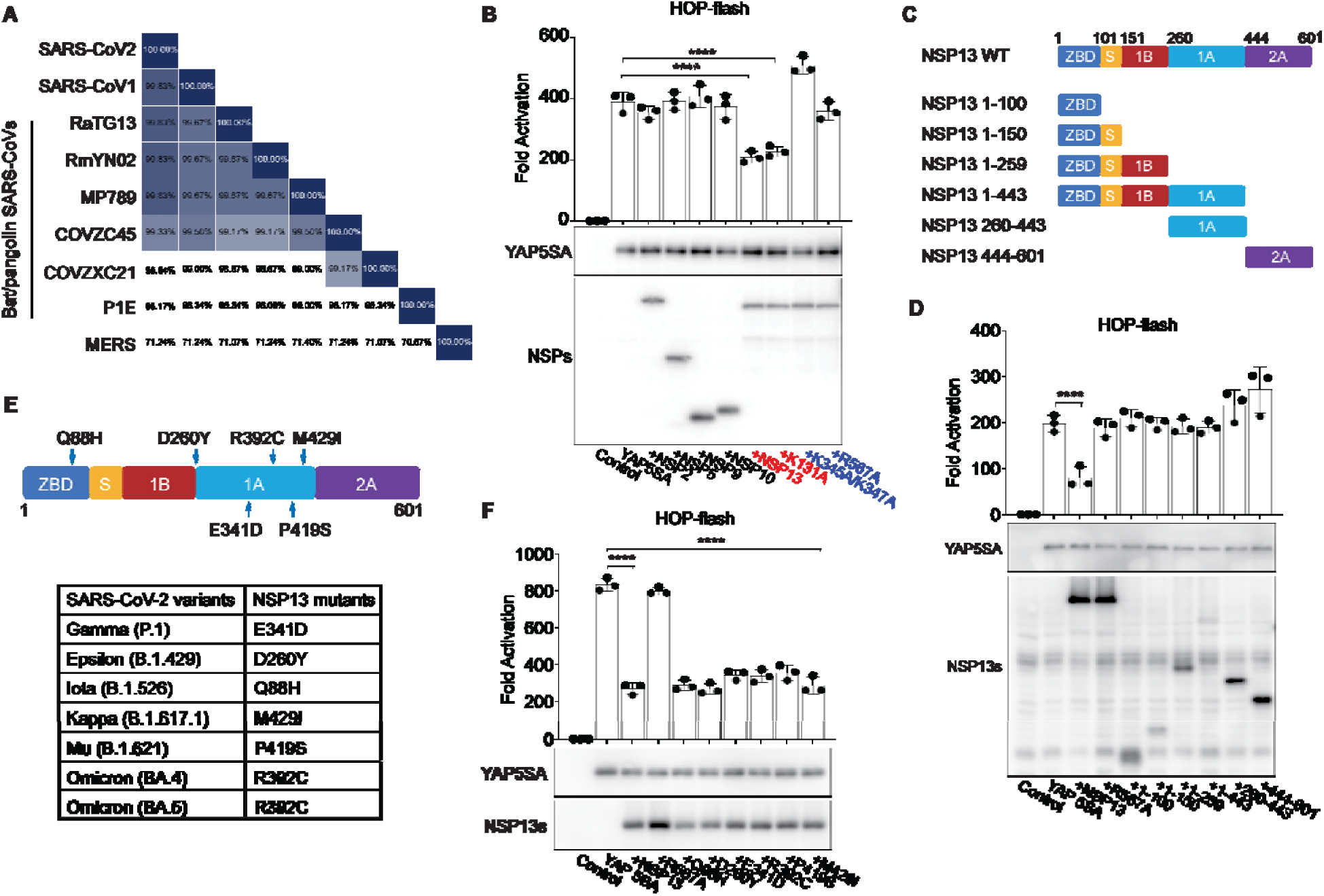
NSP13 helicase activity is required for suppressing YAP activity. (**A**) Conserved amino acid sequences of NSP13 among coronaviruses. (**B**) SARS-CoV-2 NSP13 mutant plasmids were constructed to examine YAP suppression mechanisms. NSP13-R567A, which loses its ATP consumption ability, did not inhibit YAP5SA transactivation, whereas NSP13 K345A/K347A, which loses its nucleic acid binding activity, mildly promoted YAP5SA transactivation. (n = 3 independent experiments; data are reported as the mean ± SD). **p < 0.01, ****p< 0.0001, one-way ANOVA. (**C**) 6 NSP13 truncations were contructed based on the NSP13 domain map. (**D**) Reporter assay: none of the truncations led to a reduction in YAP transactivation, and the NSP13 DNA binding domains 1A and 2A slightly increased YAP5SA activation, suggesting that the full-length NSP13 with helicase activity may be required for suppression of YAP transactivation. (n = 3 independent experiments; data are reported as the mean ± SD). *p< 0.05, ****p<0.0001, one-way ANOVA. (**E**) Summary of NSP13 mutants from SARS-CoV-2 variants. (**F**) HOP-flash reporter assay: NSP13 mutations did not affect suppression of YAP5SA transactivation. ****p < 0.0001, one-way ANOVA.

SARS-CoV-2 has consistently mutated over time, resulting in variants that differ from the original virus. We evaluated NSP13 mutations (*35*) (Figure 3E and Figure Supplement 3A) and observed that all mutants can suppress YAP transactivation (Figure 3F and Figure Supplement 3B).

### NSP13 interacts with TEAD4 and recruits YAP repressors

To determine the molecular mechanism(s) underlying NSP13 suppression of YAP, we performed immunofluorescence (IF) analyses. Our data revealed that NSP13 localizes to the cytoplasm and nucleus, whereas most YAP5SA colocalizes with NSP13 in the nucleus three days after tamoxifen injection (Figure 4A). However, co-immunoprecipitation (co-IP) assays revealed no direct interaction between NSP13 and YAP5SA (Figure Supplement 4A). Additional assays indicated that NSP13 is associated with the transcription factor TEAD4 (Figure 4B), and both the N– and C-terminal ends of TEAD4 interact with NSP13 (Figure Supplement 4B), suggesting that NSP13 prevents YAP transactivation by competitively interacting with TEAD4. Unexpectedly, introduction of NSP13 had no effect on the YAP/TEAD4 association (Figure Supplement 4C). However, the YAP/NSP13 interaction was much stronger in the presence of TEAD4, indicating that TEAD4 is a platform to recruit YAP and NSP13 (Figure 4C). Consistent with this, IF analysis of HeLa cells and heart sections revealed that YAP5SA is restricted to the nucleus when co-expressed with either NSP13 WT or R567A, in contrast to the vector control (Figure 4D, Figure Supplement 4D). Moreover, NSP13 protein levels in cardiomyocytes accumulated following YAP5SA induction (Figure 4D, E), suggesting that nuclear NSP13/YAP/TEAD4 prevents NSP13 degradation. Given that NSP13 preferentially interacts with TEAD4 in the nucleus (Figure 4B-C, Figure Supplement 4C), we next determined whether this association was DNA dependent using multiple nucleases: Universal Nuclease (which degrades all forms of DNA and RNA), DNase I (which cleaves both single and double stranded DNA), and RNase H (which selectively cleaves the RNA strand in RNA/DNA hybrids). These treatments did not disrupt the NSP13/TEAD4 interaction (Figure Supplement 4E), revealing that their binding is not dependent on nucleic acids.

**Figure 4.**
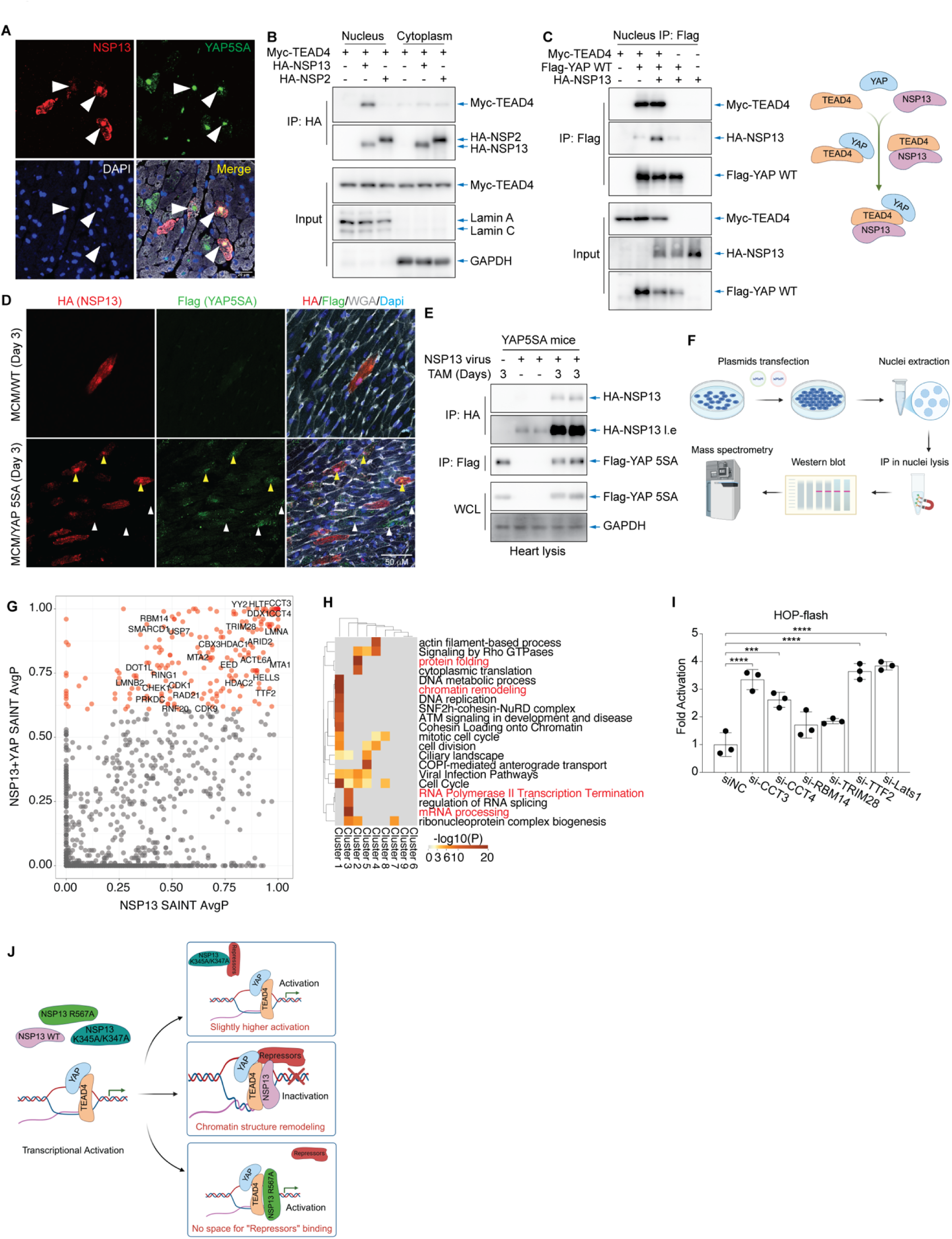
NSP13 inactivates the YAP/TEAD4 complex by recruiting YAP repressors. (**A**) Immunofluorescence imaging showing that NSP13 colocalized with YAP5SA in cardiomyocytes of YAP5SA transgenic mice 3 days after tamoxifen injection. (**B**) Co-IP: NSP13 interacts with TEAD4, a major binding partner of YAP, in the nucleus. (**C**) Co-IP of HEK293T nuclei: NSP13 does not disrupt the interaction between YAP and TEAD4, whereas TEAD4 promotes the interaction between YAP and NSP13. (**D-E**) Immunofluorescence imaging and western blot analysis reveal that NSP13 protein levels increase after YAP5SA expression is induced in YAP5SA mouse cardiomyocytes. (**F**) Workflow of IP-MS. **(G)** IP-MS in nuclei (IP: NSP13), suggesting NSP13 interacts with proteins with or without YAP co-expression. Significance Analysis of INTeractome (SAINT), AvgP >0.6 are labeled in red. (**H**) GO analysis in subclusters of NSP13 interacting proteins (SAINT, AvgP >0.6, labelled with red in Figure S4C). (**I**) HOP-flash reporter assay: endogenous YAP activity is increased after the siRNA knockdown of CCT3 and TTF2 in HeLa cells. ***p< 0.001, ****p< 0.0001, one-way ANOVA. (**J**) Working model for NSP13 regulation of YAP/TEAD.

Given that NSP13 forms a complex with YAP and TEAD4, we hypothesized that it recruits nuclear repressors to inhibit YAP transactivation. To investigate this, we performed immunoprecipitation followed by mass spectrometry (IP-MS) to identify NSP13-interacting proteins in the presence or absence of YAP in HEK293T cell nuclei (Figure 4F). Our analyses identified hundreds of candidate proteins that interact with NSP13 in the nucleus (Figure 4G), which were further analyzed by STRING-based network analysis (Figure Supplement 5A). Gene Ontology (GO) analysis indicated that the largest clusters are involved in RNA polymerase II transcription termination, chromatin remodeling, and protein folding (Figure 4H). To validate these candidate interactors, we evaluated several proteins—CCT3, SMARCD1, EIF4A1, LMNA, TTF2, and YY2—and conducted co-IP assays. Whereas TEAD4 strongly interacts with NSP13 (Figure Supplement 5B), the candidate repressors exhibited relatively weak binding to NSP13, suggesting that NSP13 associates with these proteins indirectly, potentially through a larger multiprotein complex or via chromatin-dependent interactions (Figure 4J).

Next, we evaluated the functional consequence of siRNA knockdown of selected candidates in YAP transactivation (Figure Supplement 5C). Reporter assays revealed that knockdown of CCT3 and TTF2 increased endogenous YAP transactivation in HeLa cells (Figure 4I and Figure Supplement 5D). To confirm the function of CCT3, we performed quantitative PCR analysis of classical YAP target genes and observed increased expression of *Ctgf* and *Cyr61* following *Cct3* knockdown. These findings support a model where NSP13 suppresses YAP activity by recruiting suppressors to the YAP/TEAD4 complex.

## Discussion

In this study, we observed that SARS-CoV-2 infection in human lung and hiPSC-CMs reduces YAP transcriptional activity, wherein the SARS-CoV-2 helicase NSP13 significantly inhibited YAP activity. Mechanistically, NSP13 directly interacts with TEAD4 to form a YAP/TEAD4/NSP13 complex in the nucleus, which recruits suppressors such as TTF2 and CCT3 to repress YAP-TEAD transcriptional activity (Figure 4J).

Our data reveal a helicase dependent inhibition of YAP by NSP13, where the K131, K345/K347, and R567 NSP13 residues are required. Based on published structural and biochemical studies, each of these residues uniquely supports helicase function: Substituting K131 with alanine (K131A) severely reduces helicase efficiency; K345/K347 are key DNA binding residues as mutating both (K345A/K347A) abolishes DNA binding; mutation of the ATP hydrolysis residue R567 (R567A) disables DNA unwinding. As illustrated in Fig. 4J, NSP13 must bind DNA and hydrolyze ATP to unwind nucleic acids. This helicase dependent process likely enables NSP13 to remodel chromatin by binding TEAD and organizing YAP repressors at the YAP/TEAD complex to prevent YAP/TEAD transactivation. In support of this mechanism, the K345A/K347A mutant, unable to anchor to DNA, fails to repress YAP as YAP driven transcription is slightly increased in this mutant (Figure 3B). Likewise, the ATPase dead R567A can bind DNA but does not unwind and remodel chromatin to recruit YAP repressors, resulting in a loss of YAP suppression (Figure 3B and 3F). Our model demonstrates that both DNA binding and ATP dependent unwinding are essential for NSP13 to suppress YAP transcriptional activity.

TEAD transcription factors, the major partners of YAP, are the primary nuclear effectors of Hippo-YAP signaling and play critical roles in cancer development (*11, 41*). We observed that NSP13 interacts with TEAD4, but does not disrupt the YAP/TEAD4 interaction, suggesting a novel regulatory mechanism for TEAD. IP-MS revealed that NSP13-interacting proteins such as TTF2 and CCT3 suppress the YAP-TEAD4 complex. TTF2, a SWI2/SNF2 family member, facilitates the removal of RNA polymerase II from the DNA template via ATP hydrolysis (*42, 43*). Importantly, termination of transcription complex elongation by TTF2 appears minimally affected by template position (*43*). CCT3 is a component of the chaperonin-containing T-complex, a molecular chaperone complex that promotes protein folding following ATP hydrolysis. Previous reports revealed that the liver cancer biomarker CCT3 (*44*) positively regulates YAP protein stability. We found that CCT3 interacts with NSP13 and inhibits YAP transactivation. Moreover, CCT3 knockdown increases YAP target gene expression, suggesting a context-dependent role of CCT3 in YAP regulation. While Co-IP assays detected weak interactions between NSP13 and CCT3 or TTF2, a strong NSP13/TEAD4 interaction was evident. These data suggest that NSP13 associates with CCT3 and TTF2 indirectly as part of a multiprotein complex in a chromatin structure-dependent manner, although further study is required to elucidate these mechanisms.

In conclusion, our study reveals a novel function of NSP13 in suppressing the YAP/TEAD transcriptional complex. These findings advance our understanding of the pathological effects of SARS-CoV-2 on host cells. Given that NSP13 directly interacts with TEAD4 and strongly suppresses YAP transactivation, this work provides a new avenue for modulating YAP activity as a potential treatment for YAP-driven diseases.

## Materials and methods

### Mice

We used αMyHC-MerCreMer mice, wild-type (WT) mice, and αMyHC-MerCreMer; YAP5SA mice (*24*). Mice had a mixed genetic background of C57BL/6 and 129SV. The AAV9 virus (a total of 5 × 10^11^ viral genomes, 120 μl total volume) was delivered by retro-orbital injection two weeks before tamoxifen injection. For the survival experiment in Figure 2, two low doses of tamoxifen (10 ug/g) were administered to 6-week-old mice by intraperitoneal injection. The mouse cardiac function was evaluated by echocardiography at day 4 and day 8 post-tamoxifen injection, and all the surviving mice were sacrificed at day 21. For the experiments in Figure 4, two tamoxifen doses were injected (Figure 4A, 10 ug/g; Figure 4D-E, 50 ug/g), and mouse hearts were collected at day three post-tamoxifen injection.

### RNA-seq analysis

To analyze lung samples from COVID-19 patients, snRNA sequencing data of 7 control lungs and 35 COVID-19 lungs from GSE171668 (*25*) and GSE171524 (*26*) were downloaded and analyzed using the Seurat v4 software suite (*45*). A total of 223,106 nuclei were used in the analysis. Each sample was normalized and batch corrected using SCTransformation. Mitochondrial percentage was used to regress out technical variability between batches, and we used harmony (*46*) integration to remove technical variation among samples. The YAP score was evaluated using 38 canonical YAP target genes. For analyzing iPSC-CMs infected with SARS-CoV-2, bulk RNA-seq data were obtained from Perez-Bermejo et al. (*23*) With this dataset, we re-analyzed and evaluated the average expression of 302 cardiomyocyte-specific YAP target genes (*24*).

### Expression plasmids

Expression plasmids encoding HA-tagged WT, mutant, or truncated NSP13 were generated by polymerase chain reaction (PCR) and subcloned into pXF4H (N-terminal HA tag) derived from pRK5 (Genetech). Myc-tagged WT or truncated TEAD4, flag-tagged WT YAP, flag-tagged YAP5SA, and Myc-tagged NSP13 interacting proteins, including CCT3, SMARCD1, EIF4A1, LMNA, TTF2, and YY2, were generated by PCR and subcloned into a pcDNA3 backbone. HA-tagged WT LATS2, LATS2 KR, and pRL-TK_Luc were gifts from Dr. Pinglong Xu’s lab. Reporters of HOP-flash (#83467) and 8xGTIIC-luciferase (#34615) were purchased from Addgene. The NSP plasmids used in YAP transactivation screening experiments were gifts from Dr. Nevan J. Krogan’s lab. All plasmids were confirmed by performing DNA sequencing.

### Cell culture and transfection

HEK293T and HeLa cells were cultured in Dulbecco’s Modified Eagle Medium with 10% fetal bovine serum. Lipofectamine™ 2000 (ThermoFisher) or Lipofectamine™ 3000 (ThermoFisher) reagents were used for plasmid transfection. LipofectAmine RNAiMAX (ThermoFisher) was used for siRNA transfection.

### Immunoprecipitation-mass spectrometry, database search, and data analysis

HEK293T cells were plated on a 150 mm tissue culture dish using 10% FBS DMEM 24 hours post-plating. Cells were transfected with Flag-YAP1, HA-NSP13, or Flag-YAP1+HA-NSP13 by Lipofectamine 3000 using standard manufacturer’s protocols. Each experimental group was performed in duplicate to ensure reproducibility and reliability. 24 hours post-transfection, cells were collected in a hypotonic lysis buffer (5 mM NaCl, 20 mM HEPES, pH 7.5, 0.4% NP-40) supplemented with protease inhibitor (Roche) and phosphatase inhibitor (Roche). Cell suspensions were drawn into a 3 ml syringe using a 25G needle and expelled in one rapid stroke into a new tube. After three rounds of lysis, the suspension was centrifuged at 600 x g 4 °C for 10 minutes to pellet nuclei. The supernatant was collected in a different tube, and the nuclear fraction was washed in the buffer twice. After the final wash, nuclei were resuspended in 20 mM HEPS, 150 mM NaCl, 2.5 mM MgCl2, 0.1% NP 40, and nuclear fractions were sonicated using Bioruptor® Pico for 10 cycles of 30 sec on and 30 sec off. Subsequently, lysate was centrifuged for 20 mins at 2 x 1000 g at 4°C. The supernatant was incubated with HA magnetic beads (ThermoFisher) for two hours at 4°C with continuous rotation. Antibody and protein-bound beads were washed thrice for 15 minutes in lysis buffer at 4 °C with constant rotation. Washed beads were boiled in 40 μL of 1X NUPAGE® LDS sample buffer (Invitrogen) and subjected to SDS-PAGE (NuPAGE 10% Bis-Tris Gel, Invitrogen). Eluted proteins were visualized with Coomassie Brilliant blue stain and excised into gel pieces according to molecular size. Individual gel pieces were destained and subject to in-gel digestion using trypsin (GenDepot T9600). Tryptic peptides were resuspended in 10 μl of loading solution (5% methanol containing 0.1% formic acid) and subjected to nanoflow LC-MS/MS analysis with a nano-LC 1000 system (Thermo Scientific) coupled to Orbitrap Elite (Thermo Scientific) mass spectrometer. Peptides were loaded onto a Reprosil-Pur Basic C18 (1.9 µm, Dr.Maisch GmbH, Germany) precolumn of 2 cm X 100 µm size. The precolumn was switched in-line with an in-house 50 mm x 150 µm analytical column packed with Reprosil-Pur Basic C18 equilibrated in 0.1% formic acid/water. Peptides were eluted using a 75 min discontinuous gradient of 4-26% acetonitrile/0.1% formic acid at a 700 nl/min flow rate. Eluted peptides were directly electro-sprayed into Orbitrap Elite mass spectrometer operated in the data-dependent acquisition mode acquiring fragmentation spectra of the top 25 strongest ions and under the direct control of Xcalibur software (Thermo Scietific).

MS/MS spectra were analyzed using the target-decoy mouse RefSeq database in the Proteome Discoverer 1.4 interface (Thermo Fisher), employing the Mascot algorithm (Mascot 2.4, Matrix Science). The precursor mass tolerance was set to 20 ppm, with a fragment mass tolerance of 0.5 daltons and a maximum of two allowed missed cleavages. Dynamic modifications included oxidation, protein N-terminal acetylation, and destreaking. Peptides identified from the Mascot results were validated at a 5% false discovery rate (FDR), and the iBAQ algorithm was applied to assess protein abundance, enabling comparisons of relative quantities among different proteins in the sample. The iBAQ value was determined by normalizing the summed peptide intensity by the number of theoretically observable tryptic peptides for each protein. PSM values from Proteome Discover were uploaded to the SAINT user interface on the CRAPOME website for SAINT analysis. SAINT probability and fold change were calculated using YAP IP as a user-defined control.

### Luciferase reporter assay

HEK293T or HeLa cells were transfected with WT YAP-or YAP5SA-responsive HOP-flash plasmid or 8xGTIIC-luciferase reporter, which has an open reading frame encoding firefly luciferase, along with the pRL-Luc with *Renilla* luciferase coding as the internal control for transfection and other expression vectors (NSPs), as specified. After 24 h of transfection with the indicated treatments, cells were lysed with passive lysis buffer (Promega). Luciferase assays were performed by using a dual-luciferase assay kit (Promega), and data were quantified with POLARstar Omega (BMG Labtech) and normalized to the internal *Renilla* luciferase control.

### AAV9 viral packaging

Viral vectors were used as previously described (*47*). The construct containing HA-tagged NSP13 was cloned into the pENN.AAV.cTNT, p1967-Q vector (AAV9-HA-NSP13). Empty vector–encoding green fluorescent protein was used as the control (AAV9-GFP). Both vectors were packaged into the muscle-trophic serotype AAV9 by the Intellectual and Developmental Disabilities Research Center Neuroconnectivity Core at Baylor College of Medicine. After being titered, viruses were aliquoted (1 × 10^13^ viral genome particles per tube), immediately frozen, and stored long-term at −80°C. Each aliquot was diluted in saline to make a 120-ul injection solution.

### Echocardiography

For cardiac function analysis, echocardiography was performed on a VisualSonics Vevo 2100 system with a 550-s probe. B-mode images and M-mode images were captured on a short-axis projection. Ejection fraction, fractional shortening, diameter of diastolic left ventricle, diameter of systolic left ventricle, and end-diastolic volume were calculated by using a cardiac measurement package installed in the Vevo2100 system.

### Histology and Immunofluorescence staining

Hearts were fixed in 4% paraformaldehyde overnight at 4 °C, dehydrated in serial ethanol and xylene solutions, and embedded in paraffin. For immunofluorescence staining, slides were sectioned at 7-μm intervals. For paraffin sections, samples were deparaffinized and rehydrated, treated with 3% H_2_O_2_ in EtOH and then with antigen retrieval solution (Vector Laboratories Inc., Burlingame, CA, USA), blocked with 10% donkey serum in phosphate-buffered saline, and incubated with primary antibodies. Antibodies used were rabbit anti-HA (#3724, Cell signaling) and rat anti-flag (NBP1-06712, Novus Bio-logicals). Immunofluorescence-stained images were captured on a Zeiss LSM 780 NLO Confocal/2-hoton microscope.

### Statistical Analysis

For the data in our manuscript, the specific statistical test used is presented in the figure legend. In snRNA-seq analyses, Yap score differences between AT1 from Control and COVID-19 patients were identified via the Wilcoxon test. For analyzing iPSC-CMs infected with SARS-CoV-2, the expression score of cardiomyocyte YAP targets was evaluated by Student’s t-test. Mouse survival rates in Figure 2D were analyzed by the log-rank (Mantel-Cox) test. Statistical significances were evaluated using three-way ANOVA and Šídák’s multiple comparisons test for other cardiac data, including EF, FS, and systolic LV diameter. For reporter assays in cells and heart/body ratio in Figure S2E, the statistical significance of the observed differences in mean was evaluated using a one-way or two-way ANOVA and the post hoc Tukey’s multiple comparisons test. P values less than 0.05 were considered statistically significant (*p< 0.05, **p< 0.01, ***p< 0.001, ****p< 0.0001).

## Supporting information

Supporting data values

WB raw figure

sp1 IP-MS raw data

sp2 IP-MS analysis

sp3 STRING analysis

## Acknowledgments

We are grateful to Drs. Nevan J. Krogan and Pinglong Xu for the plasmid gifts. Dr. Hongxin Guan assisted with NSP13 mutant structure analysis. Rebecca Bartow, PhD, of the Department of Scientific Publications at The Texas Heart Institute, provided editorial support.

## Funding

National Institutes of Health grant HL 127717 (JFM).

National Institutes of Health grant HL 130804 (JFM).

National Institutes of Health grant HL 118761 (JFM).

Vivian L. Smith Foundation (JFM).

2020 COVID-19 RFA (JFM).

AHA postdoctoral fellowship 903411 (FM).

AHA postdoctoral fellowship 903651 (RL).

K99 HL169742 (JS)

Don McGill Gene Editing Laboratory of The Texas Heart Institute (XL).

## Author contributions

Conceptualization: FM, JFM

Methodology: JK, JS, XL

Investigation: FM, C-RT, JW, YS, VD, BX, RL, SL

Supervision: JFM

Writing – original draft: FM

Writing – review & editing: FM, JFM

## Data and materials availability

All data are available in the main text or the supplementary materials.

## Supplemental files

**figure supplement 1.**
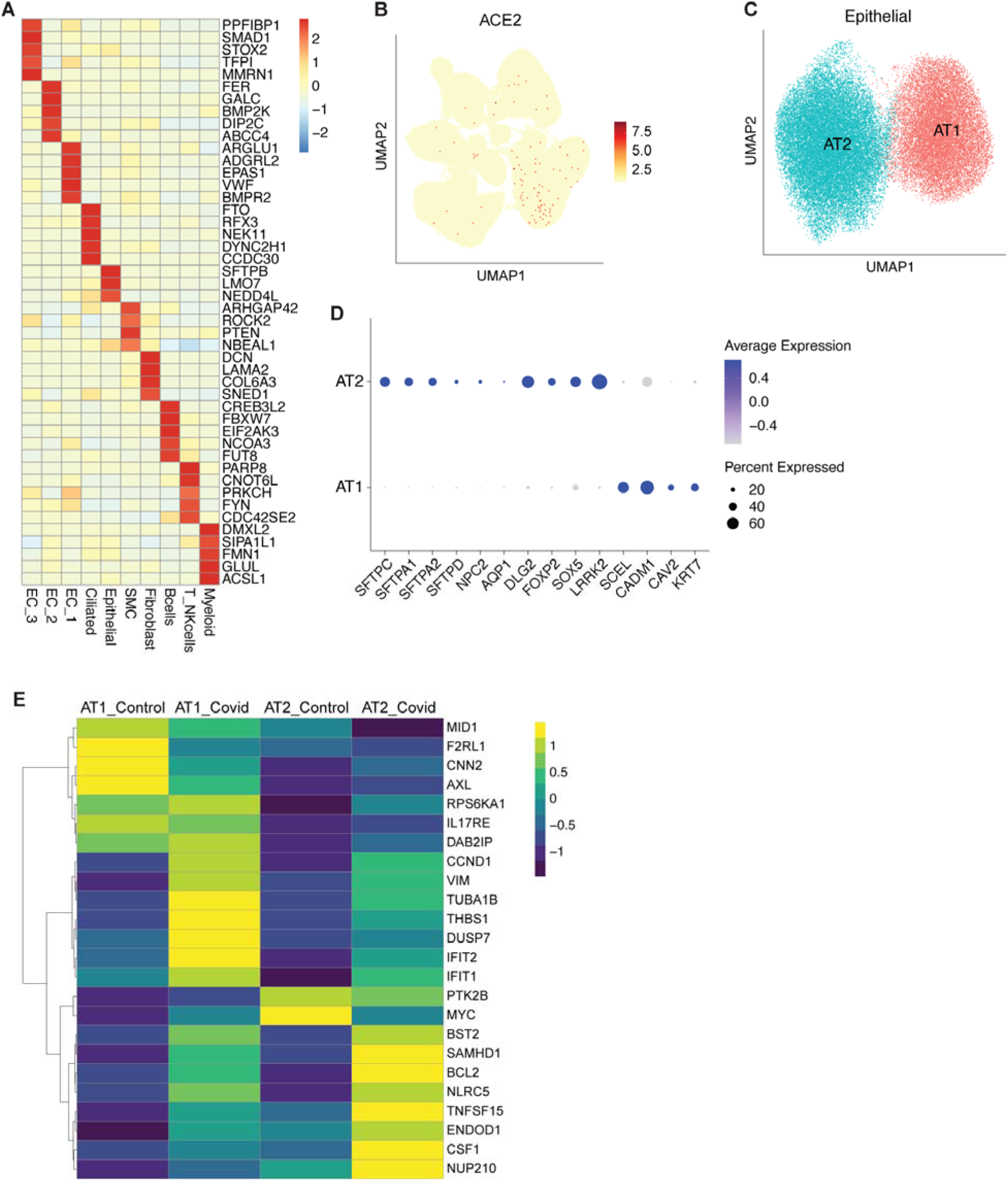
SARS-CoV-2 infection suppresses YAP activity *in vivo* and *in vitro*. (**A**) Heatmap showing the most highly expressed markers for each cell type in human lung samples. (**B**) Uniform manifold approximation and projection (UMAP) visualization showing ACE2 expression in all cell types. (**C**) UMAP of epithelial cell subclusters. (**D**) Dot plot showing the relative expression of alveolar type 1 (AT1) and alveolar type 2 (AT2) markers in human lung samples. (**E**) Expression of Yap targets genes involved in innate immune response or regulation in AT1 and AT2 cells from control and COVID-positive lung samples.

**figure supplement 2.**
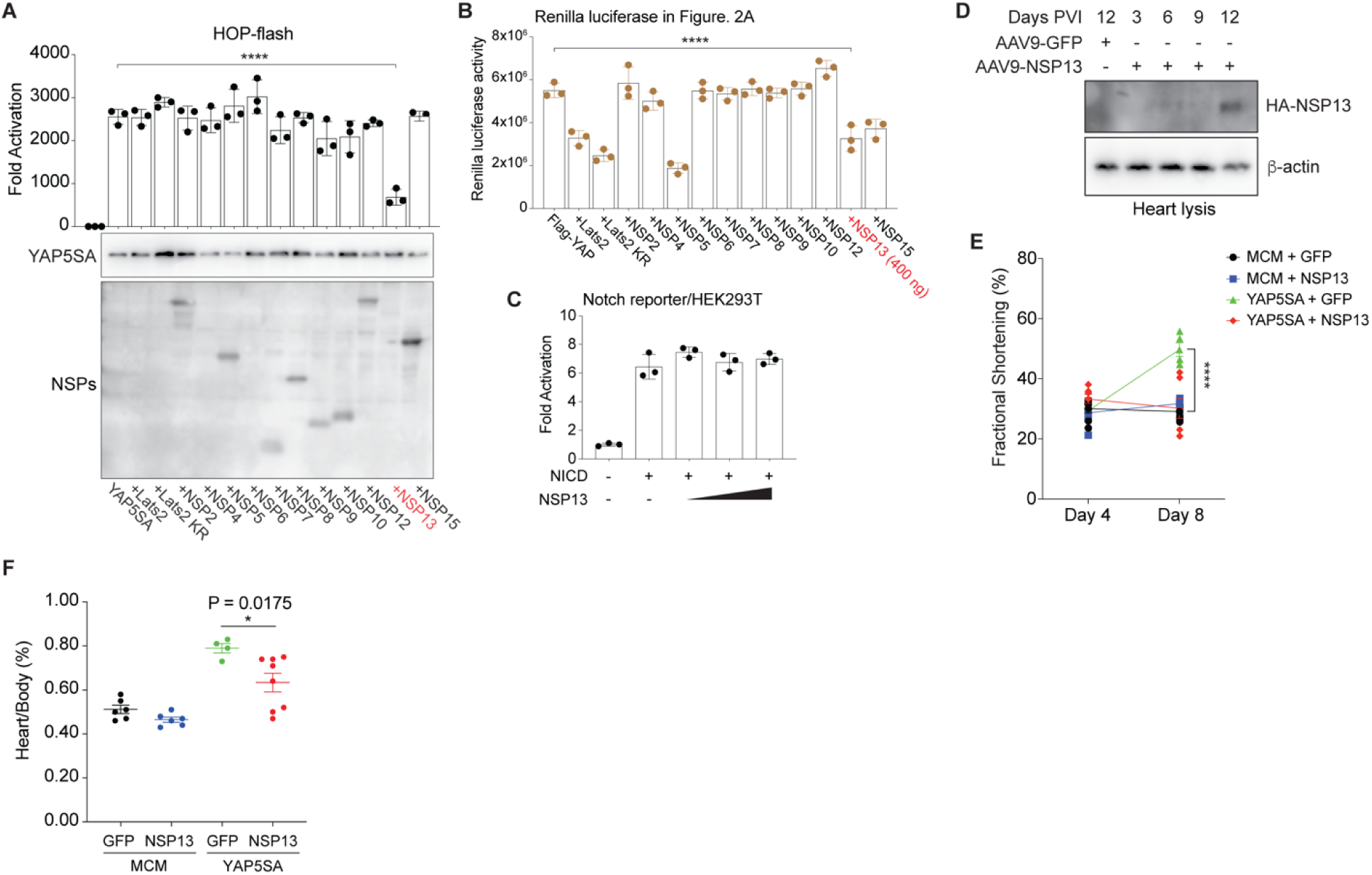
NSP13 inhibits YAP transactivation *in vitro* and *in vivo*. **(A)** Reporter assay (HOP-flash) results indicating that NSP13 inhibited YAP5SA transactivation at low protein expression levels when compared with other NSPs. (n = 3 independent experiments; data are presented as mean ± SD). ****p<0.0001, one-way ANOVA. **(B)** Raw reads of Renilla luciferase in Figure 2A. **(C)** Reporter assay (Notch reporter) results showing that NSP13 can not suppress NICD activation. (n = 3 independent experiments; data are presented as mean ± SD). (**D**) NSP13 expression at different time points in mouse hearts after AAV9-HA-NSP13 virus injection. (**E**) Fractional shortening was increased in YAP5SA mice at day 8 after tamoxifen injection (10 ug/g x2). This increase was reversed by NSP13 expression. ****p<0.0001, three-way ANOVA. (**F**) The overgrowth of the heart in YAP5SA mice, as evidenced by heart/body ratio, was reversed by NSP13 expression. *p=0.0175, one-way ANOVA.

**figure supplement 3.**
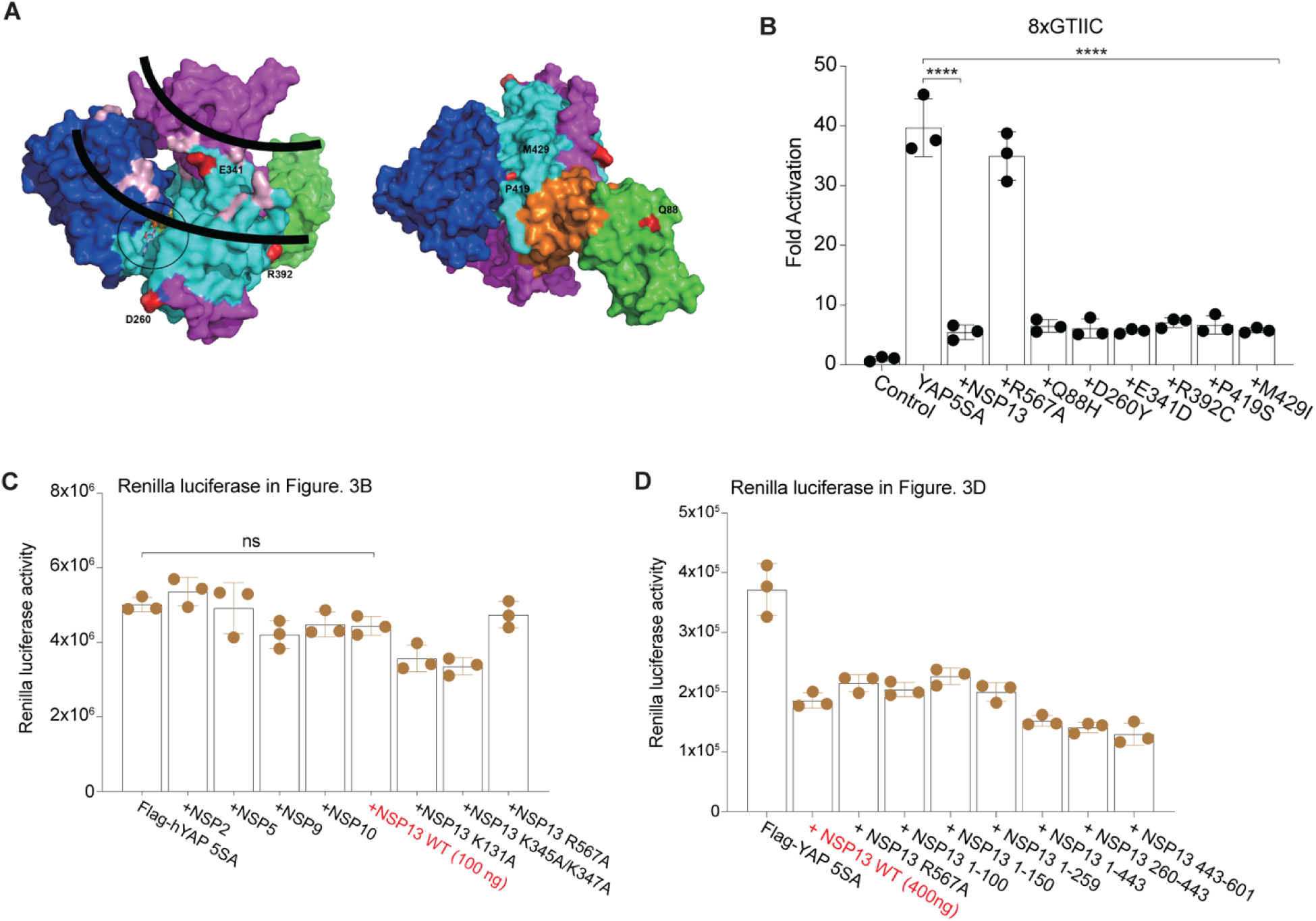
NSP13 helicase activity is required for suppressing YAP activity. (**A**) The location of the NSP13 mutations from SARS-CoV2 variants on the NSP13 protein structure model (PDB: 7NN0). (**B**) Reporter assay (8xGTIIC) results indicating that NSP13 mutations did not affect its suppression on YAP5SA transactivation. ****p<0.0001, one-way ANOVA. (**C**) Raw reads of Renilla luciferase in Figure 3B. (**D**) Raw reads of Renilla luciferase in Figure 3D.

**figure supplement 4.**
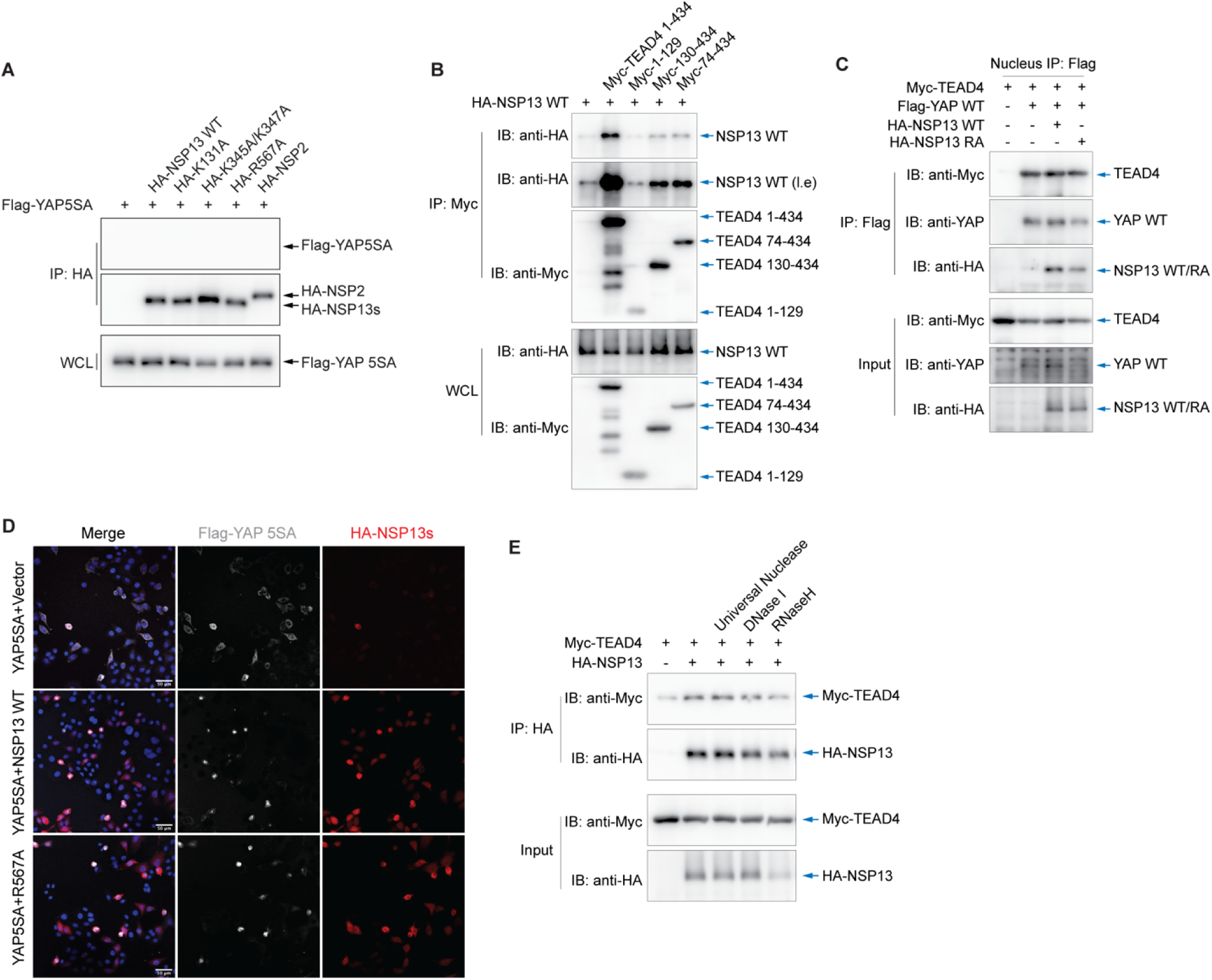
NSP13 interacts with TEAD4. (**A**) Western blot analysis showed that neither wild-type nor mutant NSP13 directly interacted with YAP5SA in HEK 293T cells. (**B**) TEAD4 domain mapping experiments showing that both the N-terminus and C-terminus are required for the interaction with NSP13. (**C**) Co-immunoprecipitation in nucleus indicated that NSP13 wild-type and R567A did not disrupt YAP and TEAD4 interaction. (**D**) Immunofluorescence imaging showing that both NSP13 WT and R567A restricted YAP5SA in cell nucleus. (**E**) Western blot analysis of co-IP showed that nucleic acid removal did not disrupt the NSP13–TEAD4 interaction.

**figure supplement 5.**
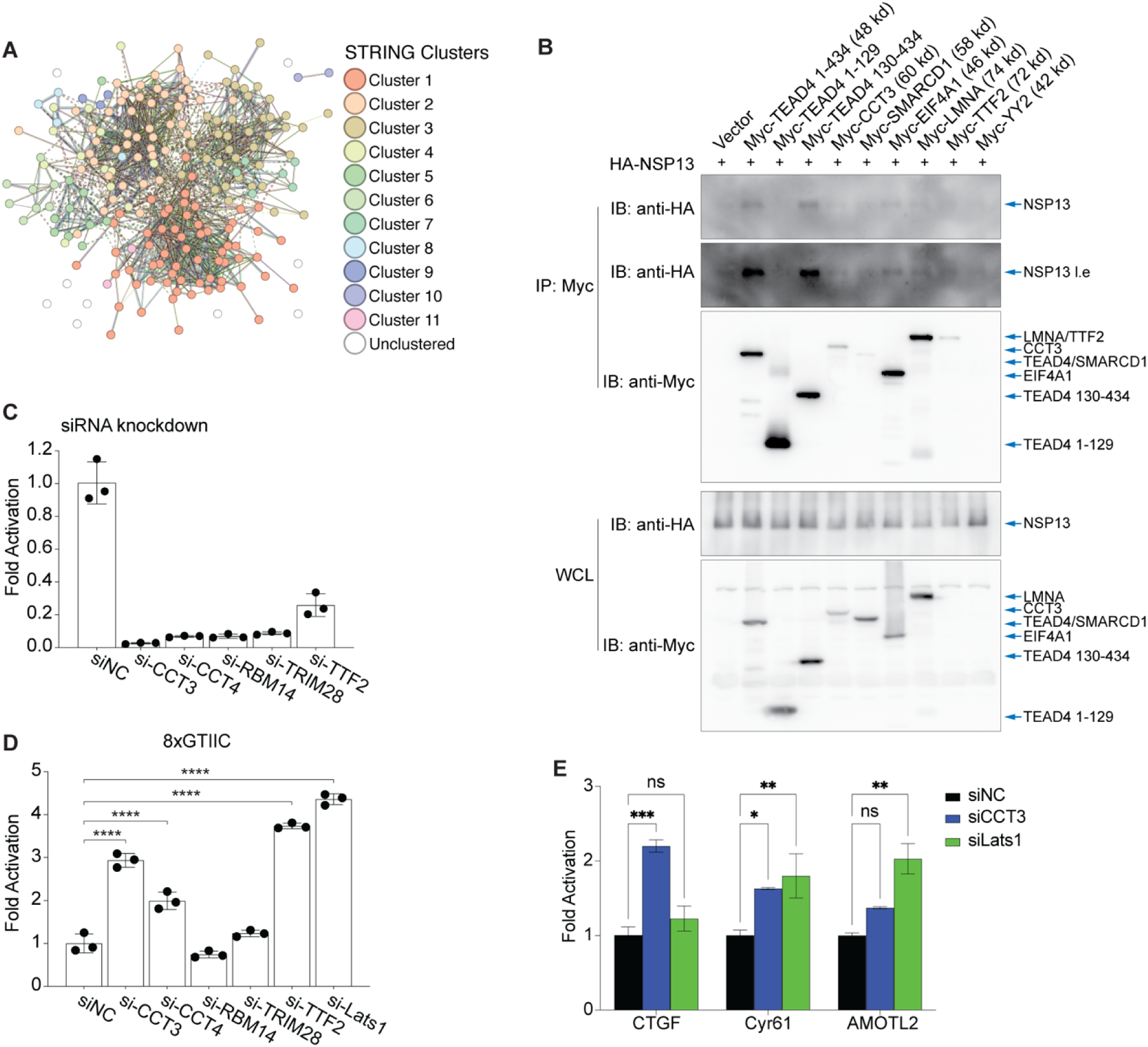
NSP13 inactivates the YAP/TEAD4 complex by recruiting YAP repressors. (**A**) Immunoprecipitation mass spectrometry in nuclei (IP: NSP13) suggesting NSP13 interacting proteins with or without YAP co-expression. Significance Analysis of INTeractome (SAINT), AvgP >0.6 were labeled with red. (**B**) co-IP assays indicated that the NSP13 interactors from mass spectrometry had weaker binding with NSP13 compared to TEAD4. (**C**) siRNA knockdown efficiency for genes in HeLa cells, determined by using quantitative polymerase chain reaction. (**D**) Reporter assay (8xGTIIC) results in HeLa cells revealed that endogenous YAP activity was increased after the siRNA-mediated knockdown of CCT3 and TTF2. ****p<0.0001, one-way ANOVA. (**E**) Quantitative PCR analysis indicated increased expression of *Ctgf* and *Cyr61* following *Cct3* knockdown. *p<0.05, **p<0.01, ***p<0.001, two-way ANOVA.

## References

1. S. Ma, Z. Meng, R. Chen, K. L. Guan, The Hippo pathway: Biology and pathophysiology. Annu. Rev. Biochem. 88, 577–604 (2019).

2. J. Wang, S. Liu, T. Heallen, J. F. Martin, The Hippo pathway in the heart: pivotal roles in development, disease, and regeneration. Nat. Rev. Cardiol 15, 672–684 (2018).

3. F. Meng, B. Xie, J. F. Martin, Targeting the Hippo pathway in heart repair. Cardiovasc. Res. 118, 2402–2414 (2022).

4. J. O. Russell, F. D. Camargo, Hippo signalling in the liver: role in development, regeneration and disease. Nat Rev Gastroenterol Hepatol 19, 297–312 (2022).

5. A. W. Hong, Z. Meng, K. L. Guan, The Hippo pathway in intestinal regeneration and disease. Nat. Rev. Gastroenterol. Hepatol. 13, 324–337 (2016).

6. I. M. Moya, G. Halder, Hippo-YAP/TAZ signalling in organ regeneration and regenerative medicine. Nat Rev Mol Cell Biol 20, 211–226 (2019).

7. Y. Zhang, H. Zhang, B. Zhao, Hippo signaling in the immune system. Trends Biochem. Sci. 43, 77–80 (2018).

8. S. Paul, S. Xie, X. Yao, A. Dey, Transcriptional Regulation of the Hippo Pathway: Current Understanding and Insights from Single-Cell Technologies. Cells 11, (2022).

9. A. Pocaterra, P. Romani, S. Dupont, YAP/TAZ functions and their regulation at a glance. J Cell Sci 133, (2020).

10. K. C. Lin, H. W. Park, K. L. Guan, Regulation of the Hippo Pathway Transcription Factor TEAD. Trends Biochem Sci 42, 862–872 (2017).

11. H. D. Huh, D. H. Kim, H. S. Jeong, H. W. Park, Regulation of TEAD transcription factors in cancer biology. Cells 8, (2019).

12. S. Wang, F. Xie, F. Chu, Z. Zhang, B. Yang, T. Dai, L. Gao, L. Wang, L. Ling, J. Jia, H. van Dam, J. Jin, L. Zhang, F. Zhou, YAP antagonizes innate antiviral immunity and is targeted for lysosomal degradation through IKKvarepsilon-mediated phosphorylation. Nat Immunol 18, 733–743 (2017).

13. Q. Zhang, F. Meng, S. Chen, S. W. Plouffe, S. Wu, S. Liu, X. Li, R. Zhou, J. Wang, B. Zhao, J. Liu, J. Qin, J. Zou, X. H. Feng, K. L. Guan, P. Xu, Hippo signalling governs cytosolic nucleic acid sensing through YAP/TAZ-mediated TBK1 blockade. Nat. Cell Biol. 19, 362–374 (2017).

14. V. S. Meli, P. K. Veerasubramanian, T. L. Downing, W. Wang, W. F. Liu, Mechanosensation to inflammation: Roles for YAP/TAZ in innate immune cells. Sci Signal 16, eadc9656 (2023).

15. S. Wang, L. Zhou, L. Ling, X. Meng, F. Chu, S. Zhang, F. Zhou, The Crosstalk Between Hippo-YAP Pathway and Innate Immunity. Front Immunol 11, 323 (2020).

16. Y. Xiao, M. C. Hill, L. Li, V. Deshmukh, T. J. Martin, J. Wang, J. F. Martin, Hippo pathway deletion in adult resting cardiac fibroblasts initiates a cell state transition with spontaneous and self-sustaining fibrosis. Genes Dev 33, 1491–1505 (2019).

17. Z. Wang, W. Lu, Y. Zhang, F. Zou, Z. Jin, T. Zhao, The Hippo Pathway and Viral Infections. Front Microbiol 10, 3033 (2019).

18. C. Ma, Y. Cong, H. Zhang, COVID-19 and the digestive system. Am. J. Gastroenterol. 115, 1003–1006 (2020).

19. F. G. De Felice, F. Tovar-Moll, J. Moll, D. P. Munoz, S. T. Ferreira, Severe acute respiratory syndrome coronavirus 2 (SARS-CoV-2) and the central nervous system. Trends Neurosci. 43, 355–357 (2020).

20. M. Nishiga, D. W. Wang, Y. Han, D. B. Lewis, J. C. Wu, COVID-19 and cardiovascular disease: from basic mechanisms to clinical perspectives. Nat. Rev. Cardiol 17, 543–558 (2020).

21. G. Garcia, Jr., A. V. Jeyachandran, Y. Wang, J. I. Irudayam, S. C. Cario, C. Sen, S. Li, Y. Li, A. Kumar, K. Nielsen-Saines, S. W. French, P. S. Shah, K. Morizono, B. N. Gomperts, A. Deb, A. Ramaiah, V. Arumugaswami, Hippo signaling pathway activation during SARS-CoV-2 infection contributes to host antiviral response. PLoS Biol 20, e3001851 (2022).

22. S. M. Pinto, Y. Subbannayya, H. Kim, L. Hagen, M. W. Gorna, A. I. Nieminen, M. Bjoras, T. Espevik, D. Kainov, R. K. Kandasamy, Multi-OMICs landscape of SARS-CoV-2-induced host responses in human lung epithelial cells. iScience 26, 105895 (2023).

23. J. A. Perez-Bermejo, S. Kang, S. J. Rockwood, C. R. Simoneau, D. A. Joy, A. C. Silva, G. N. Ramadoss, W. R. Flanigan, P. Fozouni, H. Li, P. Y. Chen, K. Nakamura, J. D. Whitman, P. J. Hanson, B. M. McManus, M. Ott, B. R. Conklin, T. C. McDevitt, SARS-CoV-2 infection of human iPSC-derived cardiac cells reflects cytopathic features in hearts of patients with COVID-19. Sci. Transl. Med. 13, (2021).

24. T. O. Monroe, M. C. Hill, Y. Morikawa, J. P. Leach, T. Heallen, S. Cao, P. H. L. Krijger, W. de Laat, X. H. T. Wehrens, G. G. Rodney, J. F. Martin, YAP partially reprograms chromatin accessibility to directly induce adult cardiogenesis in vivo. Dev. Cell 48, 765–779 e767 (2019).

25. T. M. Delorey, C. G. K. Ziegler, G. Heimberg, R. Normand, Y. Yang, A. Segerstolpe, D. Abbondanza, S. J. Fleming, A. Subramanian, D. T. Montoro, K. A. Jagadeesh, K. K. Dey, P. Sen, M. Slyper, Y. H. Pita-Juarez, D. Phillips, J. Biermann, Z. Bloom-Ackermann, N. Barkas, A. Ganna, J. Gomez, J. C. Melms, I. Katsyv, E. Normandin, P. Naderi, Y. V. Popov, S. S. Raju, S. Niezen, L. T. Tsai, K. J. Siddle, M. Sud, V. M. Tran, S. K. Vellarikkal, Y. Wang, L. Amir-Zilberstein, D. S. Atri, J. Beechem, O. R. Brook, J. Chen, P. Divakar, P. Dorceus, J. M. Engreitz, A. Essene, D. M. Fitzgerald, R. Fropf, S. Gazal, J. Gould, J. Grzyb, T. Harvey, J. Hecht, T. Hether, J. Jane-Valbuena, M. Leney-Greene, H. Ma, C. McCabe, D. E. McLoughlin, E. M. Miller, C. Muus, M. Niemi, R. Padera, L. Pan, D. Pant, C. Pe’er, J. Pfiffner-Borges, C. J. Pinto, J. Plaisted, J. Reeves, M. Ross, M. Rudy, E. H. Rueckert, M. Siciliano, A. Sturm, E. Todres, A. Waghray, S. Warren, S. Zhang, D. R. Zollinger, L. Cosimi, R. M. Gupta, N. Hacohen, H. Hibshoosh, W. Hide, A. L. Price, J. Rajagopal, P. R. Tata, S. Riedel, G. Szabo, T. L. Tickle, P. T. Ellinor, D. Hung, P. C. Sabeti, R. Novak, R. Rogers, D. E. Ingber, Z. G. Jiang, D. Juric, M. Babadi, S. L. Farhi, B. Izar, J. R. Stone, I. S. Vlachos, I. H. Solomon, O. Ashenberg, C. B. M. Porter, B. Li, A. K. Shalek, A. C. Villani, O. Rozenblatt-Rosen, A. Regev, COVID-19 tissue atlases reveal SARS-CoV-2 pathology and cellular targets. Nature 595, 107–113 (2021).

26. J. C. Melms, J. Biermann, H. Huang, Y. Wang, A. Nair, S. Tagore, I. Katsyv, A. F. Rendeiro, A. D. Amin, D. Schapiro, C. J. Frangieh, A. M. Luoma, A. Filliol, Y. Fang, H. Ravichandran, M. G. Clausi, G. A. Alba, M. Rogava, S. W. Chen, P. Ho, D. T. Montoro, A. E. Kornberg, A. S. Han, M. F. Bakhoum, N. Anandasabapathy, M. Suarez-Farinas, S. F. Bakhoum, Y. Bram, A. Borczuk, X. V. Guo, J. H. Lefkowitch, C. Marboe, S. M. Lagana, A. Del Portillo, E. J. Tsai, E. Zorn, G. S. Markowitz, R. F. Schwabe, R. E. Schwartz, O. Elemento, A. Saqi, H. Hibshoosh, J. Que, B. Izar, A molecular single-cell lung atlas of lethal COVID-19. Nature 595, 114–119 (2021).

27. M. Hoffmann, H. Kleine-Weber, S. Schroeder, N. Kruger, T. Herrler, S. Erichsen, T. S. Schiergens, G. Herrler, N. H. Wu, A. Nitsche, M. A. Muller, C. Drosten, S. Pohlmann, SARS-CoV-2 Cell Entry Depends on ACE2 and TMPRSS2 and Is Blocked by a Clinically Proven Protease Inhibitor. Cell 181, 271–280 e278 (2020).

28. I. J. Penkala, D. C. Liberti, J. Pankin, A. Sivakumar, M. M. Kremp, S. Jayachandran, J. Katzen, J. P. Leach, R. Windmueller, K. Stolz, M. P. Morley, A. Babu, S. Zhou, D. B. Frank, E. E. Morrisey, Age-dependent alveolar epithelial plasticity orchestrates lung homeostasis and regeneration. Cell Stem Cell 28, 1775–1789 e1775 (2021).

29. J. J. Gokey, J. Snowball, A. Sridharan, P. Sudha, J. A. Kitzmiller, Y. Xu, J. A. Whitsett, YAP regulates alveolar epithelial cell differentiation and AGER via NFIB/KLF5/NKX2-1. iScience 24, 102967 (2021).

30. B. Zhao, X. Wei, W. Li, R. S. Udan, Q. Yang, J. Kim, J. Xie, T. Ikenoue, J. Yu, L. Li, P. Zheng, K. Ye, A. Chinnaiyan, G. Halder, Z. C. Lai, K. L. Guan, Inactivation of YAP oncoprotein by the Hippo pathway is involved in cell contact inhibition and tissue growth control. Genes Dev 21, 2747–2761 (2007).

31. A. Li, B. Zhang, K. Zhao, Z. Yin, Y. Teng, L. Zhang, Z. Xu, K. Liang, X. Cheng, Y. Xia, SARS-CoV-2 nsp13 Restricts Episomal DNA Transcription without Affecting Chromosomal DNA. J Virol 97, e0051223 (2023).

32. L. Yan, J. Ge, L. Zheng, Y. Zhang, Y. Gao, T. Wang, Y. Huang, Y. Yang, S. Gao, M. Li, Z. Liu, H. Wang, Y. Li, Y. Chen, L. W. Guddat, Q. Wang, Z. Rao, Z. Lou, Cryo-EM structure of an extended SARS-CoV-2 replication and transcription complex reveals an intermediate state in cap synthesis. Cell 184, 184–193 e110 (2021).

33. J. Chen, B. Malone, E. Llewellyn, M. Grasso, P. M. M. Shelton, P. D. B. Olinares, K. Maruthi, E. T. Eng, H. Vatandaslar, B. T. Chait, T. M. Kapoor, S. A. Darst, E. A. Campbell, Structural basis for helicase-polymerase coupling in the SARS-CoV-2 replication-transcription complex. Cell 182, 1560–1573 e1513 (2020).

34. L. Yan, Y. Zhang, J. Ge, L. Zheng, Y. Gao, T. Wang, Z. Jia, H. Wang, Y. Huang, M. Li, Q. Wang, Z. Rao, Z. Lou, Architecture of a SARS-CoV-2 mini replication and transcription complex. Nat. Commun. 11, 5874 (2020).

35. J. A. Newman, A. Douangamath, S. Yadzani, Y. Yosaatmadja, A. Aimon, J. Brandao-Neto, L. Dunnett, T. Gorrie-Stone, R. Skyner, D. Fearon, M. Schapira, F. von Delft, O. Gileadi, Structure, mechanism and crystallographic fragment screening of the SARS-CoV-2 NSP13 helicase. Nat. Commun. 12, 4848 (2021).

36. J. Chen, Q. Wang, B. Malone, E. Llewellyn, Y. Pechersky, K. Maruthi, E. T. Eng, J. K. Perry, E. A. Campbell, D. E. Shaw, S. A. Darst, Ensemble cryo-EM reveals conformational states of the nsp13 helicase in the SARS-CoV-2 helicase replication-transcription complex. Nat Struct Mol Biol 29, 250–260 (2022).

37. B. Malone, J. Chen, Q. Wang, E. Llewellyn, Y. J. Choi, P. D. B. Olinares, X. Cao, C. Hernandez, E. T. Eng, B. T. Chait, D. E. Shaw, R. Landick, S. A. Darst, E. A. Campbell, Structural basis for backtracking by the SARS-CoV-2 replication-transcription complex. Proc Natl Acad Sci U S A 118, (2021).

38. R. Nandi, D. Bhowmik, R. Srivastava, A. Prakash, D. Kumar, Discovering potential inhibitors against SARS-CoV-2 by targeting Nsp13 Helicase. J. Biomol. Struct. Dyn. 40, 12062–12074 (2022).

39. J. Zeng, F. Weissmann, A. P. Bertolin, V. Posse, B. Canal, R. Ulferts, M. Wu, R. Harvey, S. Hussain, J. C. Milligan, C. Roustan, A. Borg, L. McCoy, L. S. Drury, S. Kjaer, J. McCauley, M. Howell, R. Beale, J. F. X. Diffley, Identifying SARS-CoV-2 antiviral compounds by screening for small molecule inhibitors of nsp13 helicase. Biochem. J. 478, 2405–2423 (2021).

40. Z. Jia, L. Yan, Z. Ren, L. Wu, J. Wang, J. Guo, L. Zheng, Z. Ming, L. Zhang, Z. Lou, Z. Rao, Delicate structural coordination of the severe acute respiratory syndrome coronavirus Nsp13 upon ATP hydrolysis. Nucleic Acids Res. 47, 6538–6550 (2019).

41. A. V. Pobbati, R. Kumar, B. P. Rubin, W. Hong, Therapeutic targeting of TEAD transcription factors in cancer. Trends Biochem Sci 48, 450–462 (2023).

42. M. Liu, Z. Xie, D. H. Price, A human RNA polymerase II transcription termination factor is a SWI2/SNF2 family member. J Biol Chem 273, 25541–25544 (1998).

43. Y. Jiang, M. Liu, C. A. Spencer, D. H. Price, Involvement of transcription termination factor 2 in mitotic repression of transcription elongation. Mol Cell 14, 375–385 (2004).

44. Y. Liu, X. Zhang, J. Lin, Y. Chen, Y. Qiao, S. Guo, Y. Yang, G. Zhu, Q. Pan, J. Wang, F. Sun, CCT3 acts upstream of YAP and TFCP2 as a potential target and tumour biomarker in liver cancer. Cell Death Dis 10, 644 (2019).

45. Y. Hao, S. Hao, E. Andersen-Nissen, W. M. Mauck, 3rd, S. Zheng, A. Butler, M. J. Lee, A. J. Wilk, C. Darby, M. Zager, P. Hoffman, M. Stoeckius, E. Papalexi, E. P. Mimitou, J. Jain, A. Srivastava, T. Stuart, L. M. Fleming, B. Yeung, A. J. Rogers, J. M. McElrath, C. A. Blish, R. Gottardo, P. Smibert, R. Satija, Integrated analysis of multimodal single-cell data. Cell 184, 3573–3587 e3529 (2021).

46. I. Korsunsky, N. Millard, J. Fan, K. Slowikowski, F. Zhang, K. Wei, Y. Baglaenko, M. Brenner, P. R. Loh, S. Raychaudhuri, Fast, sensitive and accurate integration of single-cell data with Harmony. Nat. Methods 16, 1289–1296 (2019).

47. J. P. Leach, T. Heallen, M. Zhang, M. Rahmani, Y. Morikawa, M. C. Hill, A. Segura, J. T. Willerson, J. F. Martin, Hippo pathway deficiency reverses systolic heart failure after infarction. Nature 550, 260–264 (2017).

