## Supplementary material for "SARS-CoV-2 NSP13 interacts with TEAD to suppress Hippo-YAP signaling": WB raw figure

Figure 2A

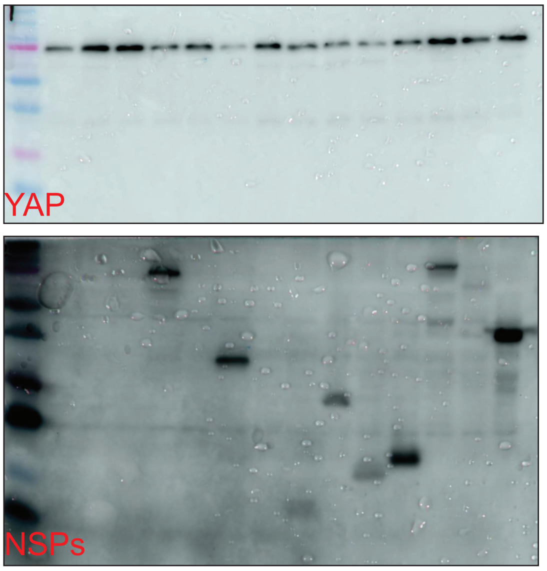


Figure 2B

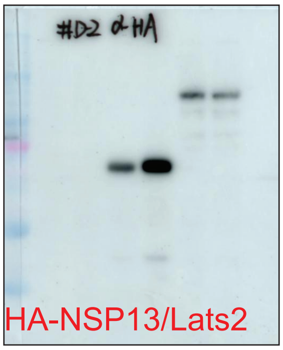


sFigure 2A

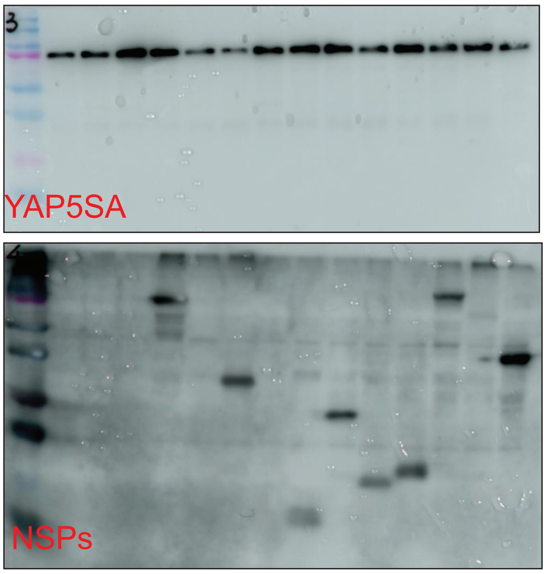


sFigure 2D

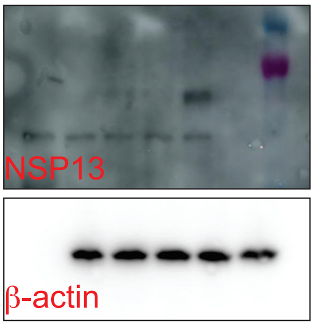


Figure 3B

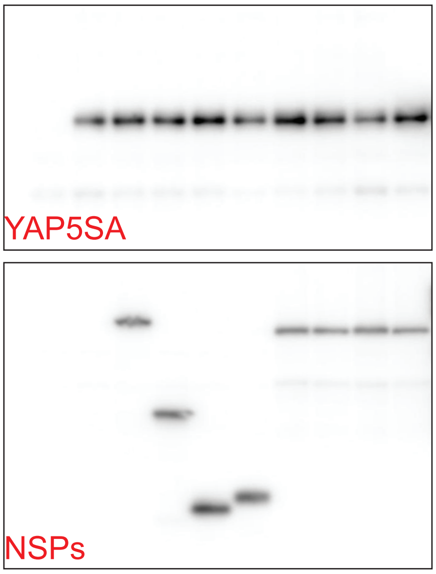


Figure 3D

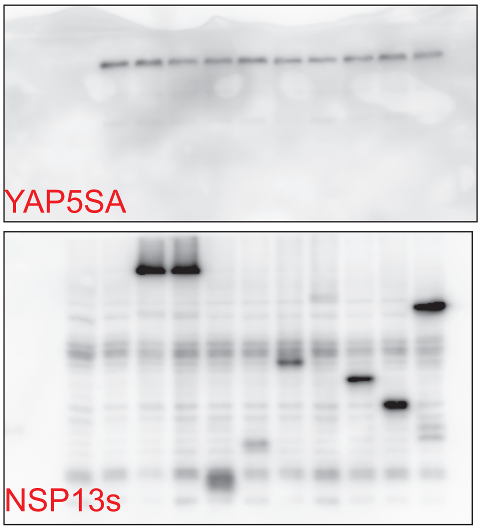


Figure 3F

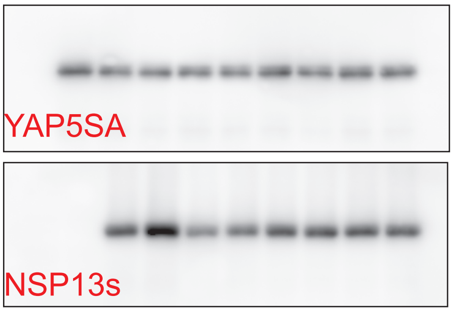

Figure 4B

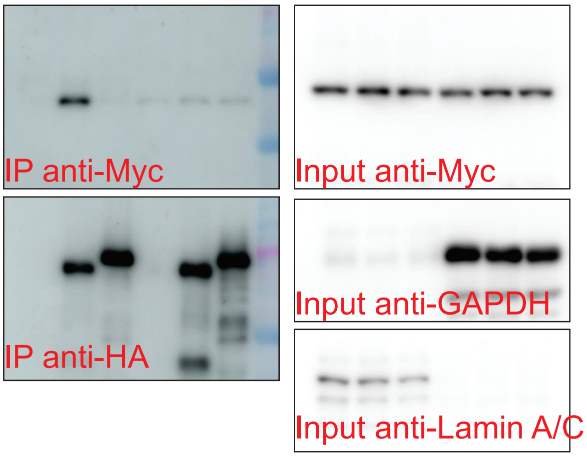


Figure 4C

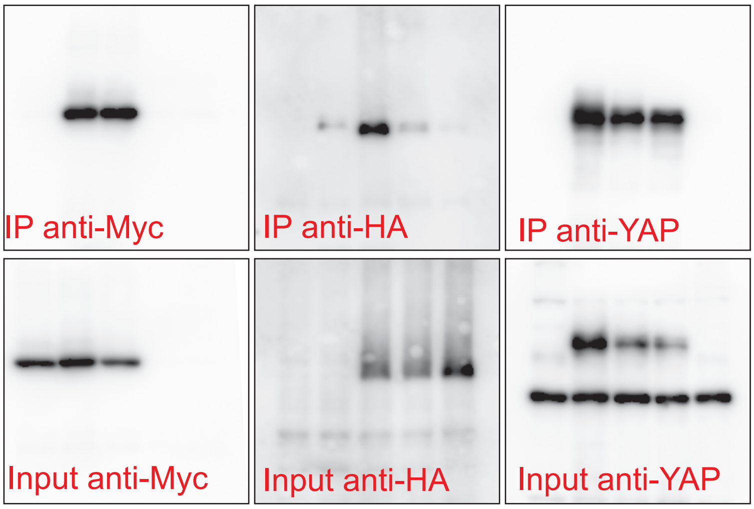


Figure 4E

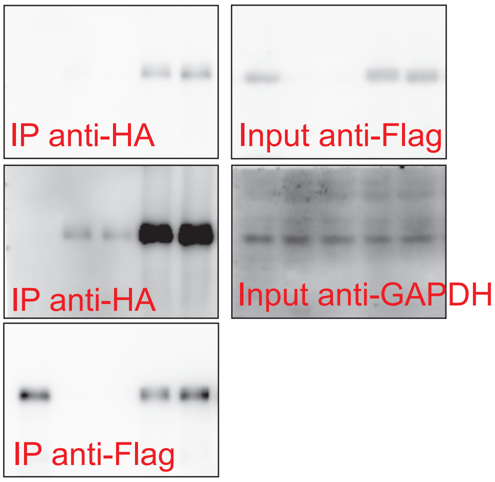


sFigure 4A

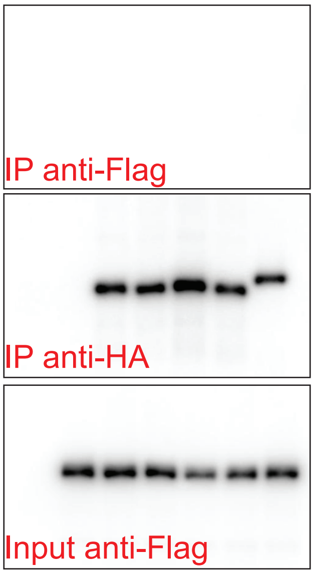


sFigure 4B

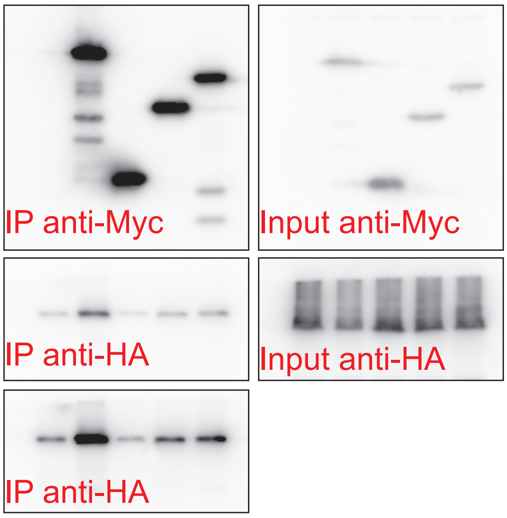


sFigure 4C

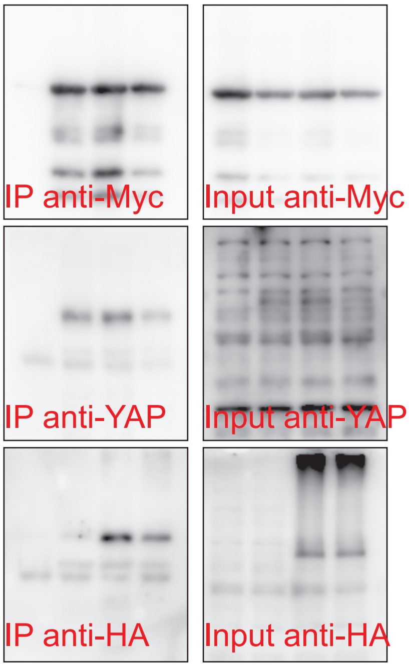


sFigure 4E

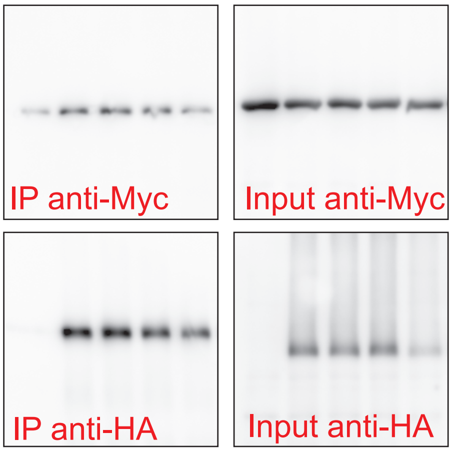

sFigure 5B

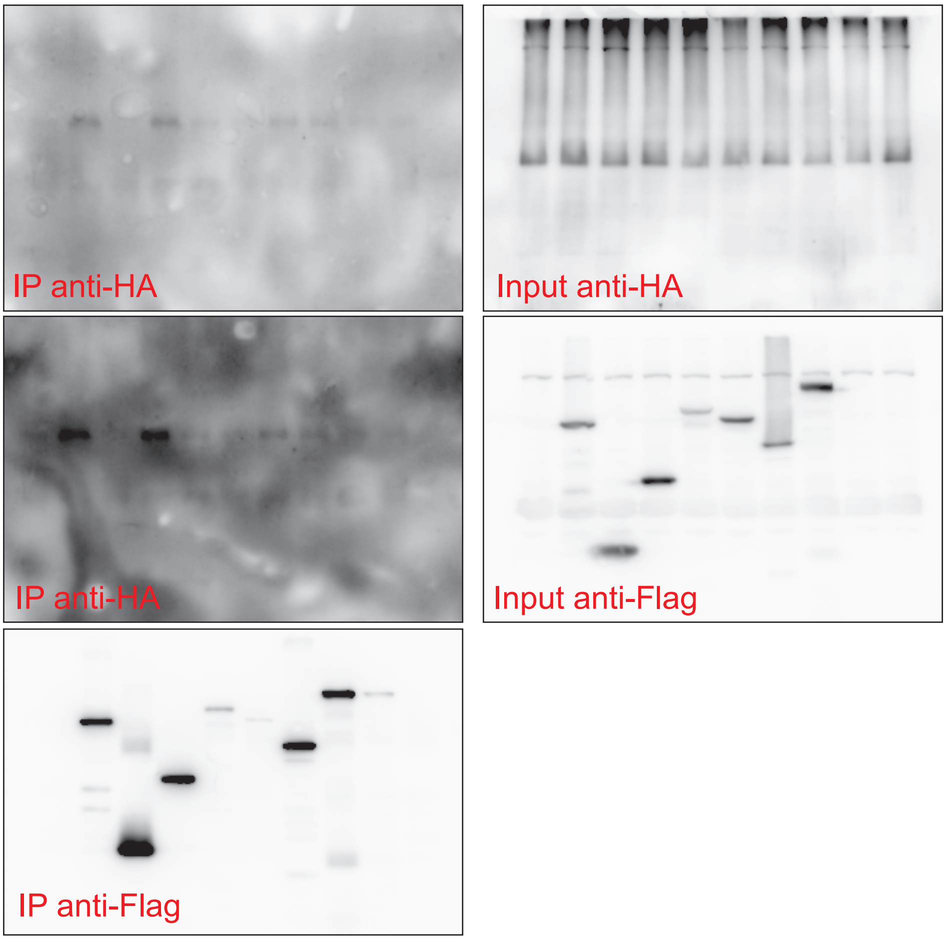
IP anti-HA
Input anti-HA
IP anti-HA
Input anti-Flag
IP anti-Flag
